# Selective activation of BK channels in small-headed dendritic spines suppresses excitatory postsynaptic potentials

**DOI:** 10.1101/2021.09.07.459293

**Authors:** Sabrina Tazerart, Maxime G. Blanchard, Soledad Miranda-Rottmann, Diana E. Mitchell, Bruno Navea Pina, Connon I. Thomas, Naomi Kamasawa, Roberto Araya

**Affiliations:** Département de Neurosciences, Université de Montréal, Montréal, Canada; The CHU Sainte-Justine Research Center; The Imaging Center and Electron Microscopy Core facility, Max Planck Florida Institute for Neuroscience, Jupiter, FL 33458, USA

**Keywords:** Synaptic transmission, pyramidal neuron, neocortex, layer 5 pyramidal neuron, BK channels, dendritic spine, two-photon (2P) uncaging, neuronal modeling, potassium channels

## Abstract

Dendritic spines are the main receptacles of excitatory information in the brain. Their particular morphology, with a small head connected to the dendrite by a slender neck, has inspired theoretical and experimental work to understand how these structural features affect the processing, storage and integration of synaptic inputs in pyramidal neurons (PNs).

The activation of glutamate receptors in spines triggers a large voltage change as well as calcium signals at the spine head. Thus, voltage-gated and calcium-activated potassium channels located in the spine head likely play a key role in synaptic transmission. Here we study the presence and function of large conductance calcium-activated potassium (BK) channels in spines from layer 5 PNs. We find that BK channels are localized to dendrites and spines regardless of their size, but their activity can only be detected in spines with small head volumes (≤ 0.09 µm^3^), which reduces the amplitude of two-photon (2P) uncaging (u) excitatory postsynaptic potentials (EPSPs) recorded at the soma. In addition, we find that calcium signals in spines with small head volumes are significantly larger than those observed in spines with larger head volumes. In accordance with our experimental data, numerical simulations predict that synaptic inputs impinging onto spines with small head volumes generate voltage responses and calcium signals within the spine head itself that are significantly larger than those observed in spines with bigger head volumes, which are sufficient to activate spine BK channels. These results show that BK channels are selectively activated in small-headed spines, suggesting a new level of dendritic spine-mediated regulation of synaptic processing, integration, and plasticity in cortical PNs.

## Introduction

Cortical pyramidal neurons (PNs) are the most abundant neurons in the cortex, with layer 5 (L5) PNs being the main output layer (Bannister, 2005; Thomson & Lamy, 2007). These neurons are the only neocortical cell type with dendrites spanning all six cortical layers, making them one of the main integrators of the cortical column. L5 PNs act as associative elements integrating sensory inputs in their proximal dendrites with information from other parts of the cortex at their distal tuft dendrites in layer 1 (Larkum, 2013).

The dendrites of L5 PNs are covered with tiny protrusions called dendritic spines (Cajal, 1888 ), which are the main recipient of excitatory inputs in the brain (Gray, 1959; Arellano *et al*., 2007b). Spines act as electrical (Koch & Poggio, 1983; Araya *et al*., 2006b; Beaulieu-Laroche & Harnett, 2018) and biochemical compartments (Yuste & Denk, 1995), which linearize the integration of subthreshold excitatory inputs in the dendrites of PNs (Araya *et al*., 2006a; Losonczy & Magee, 2006) and are the minimal functional units of synaptic plasticity (Matsuzaki *et al*., 2004; Araya, 2014; Colgan & Yasuda, 2014; Nishiyama & Yasuda, 2015; Tazerart *et al*., 2020), supporting long-term potentiation (LTP) (Lang *et al*., 2004; Matsuzaki *et al*., 2004; Tazerart *et al*., 2020) and long-term depression (LTD) (Oh *et al*., 2013) – though to be the processes directly involved in learning and memory (Makino & Malinow, 2011; Hayashi-Takagi *et al*., 2015).

The distinctive morphology of spines (Cajal, 1888 ), with a head connected to the dendrite by a slender neck (Sorra & Harris, 2000; Arellano *et al*., 2007a; Takasaki & Sabatini, 2014; Tonnesen *et al*., 2014), has inspired theoretical and experimental work suggesting that the electrotonic function of spines is to create a large voltage swing in the spine head (Miller *et al*., 1985b; Araya *et al*., 2006b; Harnett *et al*., 2012) that is sufficient to trigger the activation of voltage-gated channels, thereby modifying synaptic efficacy (Miller *et al*., 1985b).

Using two-photon (2P) uncaging of caged-glutamate onto single spines of neocortical L5 PNs or hippocampal CA1 PNs to induce synaptic activity, mimicking the synaptic release of glutamate in single dendritic spines, it has been demonstrated that dendritic spines are endowed with functional voltage-gated sodium channels that boost uncaging (u)-evoked excitatory post synaptic potentials (EPSPs) (Araya *et al*., 2007; Carter *et al*., 2012), and voltage-gated calcium channels that contribute to calcium accumulations in the spine head (Bloodgood & Sabatini, 2007). Interestingly, these channels can be activated when the 2P-uncaging of glutamate is induced right next to the spine head but not when uncaging is performed next to the parent dendrite (Araya *et al*., 2007; Bloodgood *et al*., 2009), highlighting the ability of spines to act as electrical compartments.

Furthermore, the presence of voltage-gated potassium channels in the spine can contribute to spine EPSP repolarization and, together with passive mechanisms, may account for the attenuation of synaptic potentials through the spine neck (Araya *et al*., 2006b; Araya *et al*., 2014). In fact, experiments in hippocampal and cortical PNs have shown that spines are endowed with voltage-gated A-type potassium channel Kv4.2. (Burkhalter *et al*., 2006; Kim *et al*., 2007) which affect EPSP amplitudes and the induction of LTP (Kim *et al*., 2007). In addition, small conductance calcium-activated potassium channels (SK channels) have been shown to be expressed in spines from hippocampal CA1 PNs, influencing NMDA-dependent calcium signals at the spine head, synaptic transmission, plasticity and learning and memory (Ngo-Anh *et al*., 2005; Hammond *et al*., 2006; Allen *et al*., 2011). Recently, it has been shown in L5 PNs that the activation of SK channels by backpropagating action potentials (bAPs) suppresses EPSPs and influences the induction of spike timing dependent plasticity (STDP) (Jones *et al*., 2017).

The activation of glutamate receptors in spines triggers a large voltage change (Araya *et al*., 2007; Harnett *et al*., 2012) capable of releasing magnesium block from NMDA receptors, and the opening of voltage-gated calcium channels, which leads to the accumulation of calcium at the spine head (Yuste & Denk, 1995; Bloodgood & Sabatini, 2007; Araya, 2014). Thus, voltage-gated and calcium-activated potassium channels are good candidates to be located in the spine and contribute to synaptic transmission.

Large conductance calcium-activated potassium channels (BK channels), also known as Slo1, belong to the family of Slo channels, which are comprised of 4-pore forming alpha subunits (Contreras *et al*., 2013; Latorre *et al*., 2017). A BK channel increases its open probability with large intracellular changes in calcium concentration and membrane voltages *via* modular voltage sensors and calcium binding sites (Latorre *et al*., 1989; Horrigan & Aldrich, 2002; Latorre *et al*., 2017). BK channels are ubiquitously expressed in mammals (Latorre *et al*., 1989; McManus, 1991), where they can associate with modulatory auxiliary β and γ subunits (Latorre *et al*., 2017). *In situ* hybridization and immunofluorescence studies have shown that BK channels are expressed in hippocampal mossy fiber terminals (Knaus *et al*., 1996; Misonou *et al*., 2006; Sailer *et al*., 2006). In addition, electron microscopy studies in hippocampal CA1 PNs excitatory synapses have shown the presence of BK channels at pre and postsynaptic terminals (Hu *et al*., 2001; Sailer *et al*., 2006). In L5 PNs, immunohistochemistry and electrophysiological recordings have shown that BK channels are homogeneously distributed on the soma and apical dendrites (Benhassine & Berger, 2005, 2009). The effect of BK channels in L5 PN output generation during physiological (Poolos & Johnston, 1999; Bock & Stuart, 2016) and pathological conditions has been studied (Kang *et al*., 2000). More specifically, the generation of action potentials (APs) in L5 PNs has been shown to activate somatic BK channels, which repolarize the cell and control AP duration (Benhassine & Berger, 2009; Bock & Stuart, 2016) (although contradicting results were reported (Kang *et al*., 2000)). Similar changes in AP duration have also been reported in hippocampal PNs (Lancaster & Nicoll, 1987; Storm, 1987; Shao *et al*., 1999). In addition, it has been reported in both L5 PNs (Bock & Stuart, 2016) and CA1 PNs (Golding *et al*., 1999) that the activation of dendritic BK channels regulate dendritic calcium spike duration and consequentially AP burst firing, without affecting the amplitude of bAPs (Poolos & Johnston, 1999; Bock & Stuart, 2016).

These results suggest that BK channel activation in hippocampal and cortical L5 PNs is dependent on large depolarizations and intracellular augmentation of calcium concentration, and thus its subcellular location. Since BK channel activation depends on two biophysical parameters – large depolarization and increased intracellular calcium – we reasoned that BK channels could be located in spines and contribute to synaptic transmission in L5 PNs. Our results indicate that BK channels are expressed and localized to dendrites and spines regardless of their morphology but are selectively activated in spines with small head volumes (≤ 0.09 µm^3^) - which reduces the EPSP amplitude. Furthermore, the calcium signal in activated spines (via 2P uncaging) with small head volumes is significantly larger than those observed in spines with larger head volumes, a requirement for the activation of BK channels. Indeed, numerical simulations corroborate these experimental results, showing that synaptic-induced activation of BK channels only occurs in spines with small head volumes – where large voltage responses and calcium signals are generated. Interestingly, spines where synaptic transmission can recruit BK channels also represent the population of spines with the ability to undergo synaptic plasticity (Tazerart *et al*., 2020). These results show how EPSPs are finely tuned by both passive and active spine mechanisms, which are relevant for our understanding of the relationship of spine structure and function, and the function of spines in synaptic transmission, plasticity and ultimately the input-output properties of L5 PNs.

## Material and Methods

### Animals and ethics

C57B/6 male and female mice aged P14-P21 where used for electrophysiology experiments and Western Blot analysis. Male and female mice Tg(Thy1-EGFP)MJrs/J (RRID:IMSR_JAX:007788) or B6.Cg-Tg(Thy1-YFP)HJrs/J (RRID:IMSR_JAX:003782) in a CD-1 background aged P21-P27 were used for Immunofluorescence and serial electron microscopy. These studies were performed in compliance with experimental protocols (13-185, 15-002, 16-011,17-012, 18-011 and 19-018) approved by the *Comité de déontologie de l’expérimentation sur les animaux* (CDEA) of the University of Montreal, Montreal and protocol 2020-2634 approved by the *Comité institutionnel des bonnes pratiques animales en recherche* (CIBPAR) of The Research Center at the CHU Ste-Justine, Montreal, Quebec.

### *In-situ* hybridization

*In-situ* hybridization was performed by the Allen brain Institute (Lein *et al*., 2007). The protocol at a glance is as follows: Sagittal 25 μm cryosections from a P56 male C57B/6J mouse brain were treated with peroxidase and proteinase K and hybridized with a digoxigenin-labeled riboprobe. The colorimetric signal was developed with anti-digoxigenin HRP conjugated antibody and tyramide signal amplification (TSA) followed by binding of alkaline phosphatase (AP) and the reaction of AP with 5-bromo-4-chloro-3-indolylphosphate (BCIP), which in turn reacts with nitroblue tetrazolium (NBT) to produce a blue particulate precipitate at the sites of probe binding. The antisense sequence of the c-terminal region (2902-3323) of the kcnmb1 mouse gen transcript NM_001253358.1 was used as riboprobe. This sequence has 100% homology with 5/22 splice variants and 86% with the remaining available RefSeq for kcnma1. Sagittal image 11/20 containing visual cortex was used unprocessed (http://mouse.brain-map.org/gene/show/16304). The Allen Institute mouse reference atlas was used to identify approximate cortical areas and layers.

### Synaptoneurosome (SN) preparation

An adaptation of a previously described method was used (Villasana *et al*., 2006; Seibt *et al*., 2012) as follows: P14 mouse visual cortex was dissected, flash frozen and stored at -80°C. Individual samples (approx. 1 mg) were homogenized in 700 μL SN lysis buffer (10 mM Hepes, pH 7.4, containing proteinase inhibitor) with a 2 mL Kimble-Chase tissue grinder (Thermo Fisher K885300-0002). Using 6 loose pestle and 12 tight pestle strokes. A 150 μL aliquot of this total protein (TP) fraction was boiled in 10% SDS for 10 min and stored. The remaining fraction was centrifuged at 2000xg for 2 min at 4°C to remove cellular and nuclear debris. The supernatant was loaded into a 1 mL syringe and filtered through three layers of pre-wetted 100 μm pore nylon filter (Millipore NY1H04700). The filtrate was directly loaded into 5-μm-pore centrifugal filters (Ultrafree-CL, Millipore UFC40SV25) and centrifuged at 5000xg for 15 min in a fixed angle rotor at 4 °C. The supernatant was carefully removed, and the loose pellet resuspended in boiling SN buffer (containing 2% SDS), boiled for 10 min and store at – 80°C (SN protein fraction). Protein concentration was determined with the BCA protein assay kit (Thermo Fisher 23227) using a bovine serum albumin (BSA) standard curve.

### Western Blot

An adaptation of a previously described method was used (Miranda-Rottmann *et al*., 2010) as follows: protein samples were separated in Bis-Tris 4-12% polyacrylamide gradient gels (Thermo Fisher NP0323) in MOPS-SDS running buffer (Thermo Fisher NP0001) and transferred into a PVDV membrane using a mini trans-blot cell (BioRad 170-3930) at 200V for 1h. Ponceau red staining was performed after transfer and recorded. Membranes were blocked in 0.1% Tween20 in Phosphate buffered saline (PBS) starting block (Thermo Fisher 37538) and incubated with primary antibodies diluted in the same buffer: anti-BK 1:1,000 (Antibodies Inc AB_224953) and after striping a mix of anti-γ-tubulin 1:10,000 (Sigma-Aldrich,T5326,) and anti-PSD95 1:5,000 (Antibodies Inc AB_224953). Secondary anti-mouse antibody was used 1:20,000 (Jackson ImmuneResearch 715-035-150,) and incubated for 2h at room temperature. Signal was developed using ECL substrate (BioRad170-5060) and recorded on film. Densitometric analysis was performed with the open-source program FIJI (Schindelin *et al*., 2012) and data was plotted using Prism7 (GraphPad)

### Immunofluorescence

Anesthetized P27 mice were intracardially perfused with 4% PFA, and the brain was removed and post-fixed in 4% PFA for 2h at 4°C or alternatively in 4% PFA with 0.2% picric acid and 0.05% glutaraldehyde in 100 mM Sorensen’s phosphate buffer (PB, pH 7.4). Brains were washed in PBS and dehydrated in 30% sucrose at 4°C and mounted in OCT by freezing in a bath of 2- methylbutane at -50°C. The 40 μm cryosections were cut at -18°C and placed over glass slides treated with tissue capture pen (Electron Microscopy Sciences) and stored at -80°C. The cryosections were permeabilized and blocked in TSA blocking reagent (Perkin Elmer FP1020). Primary antibodies anti-BK 1:100 Antibodies Inc AB_224953, affinity purified custom antisera against the peptide KYVQEDRL to detect short BK, and against the peptide QEKKWFTDEPDNA to detect long BK (Miranda-Rottmann *et al*., 2010) 1:40 and anti-GFP 1:400 Rockland 600-101- 215 were incubated 24h at 4° C followed by secondary antibodies: Alexa 568 anti-mouse 1:200 (Thermo Fisher A10037), Alexa 568 anti-Rabbit (Thermo Fisher A10042), and Alexa 488 anti-goat 1:200 (Thermo Fisher A11055) for 2h at room temperature (RT) in TSA blocking reagent. Washes were done in PBS. The samples were mounted in ProLong Diamond Antifade Mountant with DAPI (Thermo Fisher P36971) or Vectashield (Vector Labs H-1200) and the basal dendrites from L5 PNs were imaged using a Zeiss LSM700 confocal microscope (IRIC bio-imaging core) or a Leica TCS SP8 laser scanning confocal microscope with DLS light sheet module (PIM core). Each optical slice was 0.7- 0.9 μm thick.

### Immuno Electron Microscopy

Anesthetized P27 mice were intracardially flushed with 25 mM PB containing 150 mM NaCl, then perfused with 4% PFA with 0.2% picric acid and 0.05% glutaraldehyde in 100 mM PB. Brains were removed and post-fixed with the same fixative for 4 hours at 4 °C, then sliced at a thickness of 500 µm. Slices were cryo-protected in 10 and 20% glycerol made in 100 mM PB for 30 min each, then left overnight in 30% glycerol at 4 °C. Slices were trimmed then plunge frozen in propane cooled to -183 °C (CPC, Leica Microsystems). Freeze substitution was then performed on slices, beginning with 1% uranyl acetate in methanol for 24 h at -90 °C, increasing 4 °C every hour to -45 °C. Slices were washed in methanol for 6 h, then infiltrated in ascending concentrations of acrylic resin for 30 h (Lowicryl HM20 mixture, Electron Microscopy Science). Samples were UV polymerized starting at -45 °C, and the temperature was increased 5 °C/h up to 20 °C, and left for 24 h. Once cured, gold sections were cut (80-90 nm thin) and collected on nickel TEM grids. Grids were placed on drops of 0.1% sodium borohydride made in 50 mM Tris Buffered Saline (TBS, pH 7.4), with 50 mM glycine for 20 minutes then placed on blocking reagent for 30 min (5% bovine serum albumin, 5% normal goat serum, 0.1 M TBS). Primary antibody mixtures contained anti-BK channel (same as those used for immunofluorescence studies) diluted 1:30 either against the peptide KYVQEDRL to detect short BK or against the peptide QEKKWFTDEPDNA to detect long BK (Miranda-Rottmann *et al*., 2010), as well as 1:100 diluted chicken anti-GFP (Abcam ab13970). Samples were incubated 40-42 h at 4 °C. After washing, samples were incubated in the secondary antibody mixture: 1:30 diluted 12 nm gold conjugated donkey anti-chicken (Jackson Immunoresearch, 703-205-155), and 1:30 diluted 6 nm gold conjugated donkey anti-rabbit (Jackson Immunoresearch, 711-195-152) for 6.5 h at room temperature (RT) in 1:10 diluted blocking buffer. Washes were done in TBS. The reaction was stabilized using 1% glutaraldehyde in TBS for 5 minutes, then rinsed in water. Sections were counter-stained with 3% aqueous uranyl acetate and lead citrate for 1 minute each, then observed using a Tecnai G2 Spirit transmission electron microscope (Thermo Fisher Scientific) at 100 kV, and images were acquired using a Veleta CCD camera (Olympus) operated by TIA software (Thermo Fisher Scientific). PSD length was measured with Fiji (Schindelin *et al*., 2012) using a segmented line tool to follow the membrane above the electron dense PSD material. Estimated spine head volume was calculated by measuring the major and minor axes of the fitted ellipse to the outer shape of the spine head demarcated with a polygon tool.

### Image processing for immunofluorescence experiments

The LSM toolbox Fiji (Schindelin *et al*., 2012) plugin was used to process the confocal images as follows: Raw images saved in LSM format were opened and optical slices of the z stack with good dendritic morphology observed in the green (GFP) channel were duplicated (to extract one slice), rotated and cropped. Red (BK) and green channels were separated with the color split function and contrast and brightness was manually adjusted to achieve clear images of similar intensity in both channels. Gaussian blur set at 1-1.5 (the same for both channels) was then applied to round the pixels. Both channels were stacked using the color merge channel function which generates a yellow color (merge) in overlapping areas of similar intensity. Scale bars were automatically drawn using the LSM bar tool function. Allen Brain Atlas *in-situ* hybridization images were downloaded in full resolution, cropped and scaled using Fiji.

### Slice preparation and solutions

Following decapitation, the brain of P14-P22 C57B/6 mice was quickly placed in an ice-cold solution made of (in mM): 222 Sucrose, 27 NaHCO3, 2.6 KCl, 1.5 NaH2PO4, 3 MgSO4 and 1 CaCl2. 300 μm thick slices from the visual cortex were then obtained using a vibrating blade (Electron Microscopy Sciences) microtome (VT 1000S, Leica) and transferred to a chamber containing artificial cerebrospinal fluid (ACSF) solution kept at 32°C containing (in mM): 126 NaCl, 26 NaHCO3, 10 Dextrose, 3 KCl, 1.15 NaH2PO4, 2 MgSO4 and 2 CaCl2. Following a 30- minute incubation, slices recovered for 1 hour at room temperature (∼20°C) before experiments were started. All solutions were oxygenated using a 95%/5% mixture of O2 and CO2 (pH: 7.4; 300mOsm).

### Electrophysiology

Recordings were obtained using a MultiClamp 700B amplifier (Axon Instruments) interfaced to a dedicated computer by a BNC-2090A data acquisition board (National Instruments). The electrophysiological signals were acquired at 10 kHz using the *PackIO* open-source software package (http://www.packio.org). Offline data analysis was performed with the MATLAB (Mathworks) *EphysViewer* package (https://github.com/apacker83/EphysViewer), IgorPro (WaveMetrics) and the *NeuroMatic* IgorPro package (http://www.neuromatic.thinkrandom.com/). Pipettes were pulled from borosilicate tubing and had resistance between 3 and 5 MΩ when filled with the pipette solution (Mitchell *et al*., 2019; Tazerart *et al*., 2019). The patch pipette solution was made of (in mM): 130 potassium gluconate, 5 KCl, 2 MgCl2, 10 HEPES, 2 MgATP, 0.3 NaGTP and 0.3-0.5% biocytin.

### Two-photon (2P) imaging and glutamate uncaging

Visually identified L5 PNs were filled with Alexa Fluor 488 (200 µM, Invitrogen) or Alexa Fluor 564 (120 µM) using a somatic whole-cell patch pipette. Cells were then imaged using a BX51WI inverted microscope (Olympus) equipped with a set of laser scanning galvo mirrors, two photomultiplier tubes with red and green channels (PMT, company) and a Chameleon ultrafast laser (Chameleon, Coherent) tuned at 720 nm for both imaging and uncaging as previously described (Mitchell *et al*., 2019). The intensity of the laser on sample was controlled using a Pockel cell (Conoptics) and set to approximately 5-8mW on sample for imaging and 25-35 mW on sample for glutamate uncaging.

Imaging and glutamate uncaging were performed using the *PrairieView* software suite (Bruker). Before starting the uncaging experiments, *4-methoxy-7-nitroindoline-caged L-glutamate* (*MNI-caged-glutamate*, 2.5 mM, Tocris Bioscience) was added to the bath solution, which was recirculated by a peristaltic pump to minimize the total bath volume (≍3 mL).

Spines were identified by loading the whole-cell patch-clamped L5 PN with Alexa Fluor 488 or Alexa Fluor 564 and monitoring its fluorescence using a wavelength of 720 nm at low laser power (5-8 mW on sample). Once a spine was identified, the uncaging spot was placed near the spine head (≍0.2 µm away from the upper edge of the spine head), in the z-plane of maximal spine fluorescence intensity, and a series of 5-10 pulses of 4 ms duration were delivered at a rate of 0.5 Hz. Voltage deflections due to the glutamate uncaging (uEPSP) were recorded from the soma in whole-cell current-clamp mode, maintaining a resting potential of -65 mV. uEPSPs are reported as the average of 5-10 pulses. From the uEPSP triggered, the peak amplitude was analyzed. Since the spine head volume we targeted could vary from 0.06 to 0.3 µm^3^, the relationship between the spine head volume and the effect of BK channel blockers on uEPSP amplitude was plotted and fitted with a plateau followed by a one phase decay curve (R = 0.71, λ = 0.09 µm^3^, GraphPad Prism 5).

### Calcium measurement

For calcium imaging experiments, visually identified L5 PNs were filled with Alexa 594 (100µM) and Fluo-4 (300 μM; Thermo Fisher) through the somatic patch clamp recording pipette. To perform sequential 2P calcium imaging and 2P uncaging of caged glutamate in selected spines at one wavelength (810 nm), we used *ruthenium-bipyridine-trimethylphosphine-caged glutamate* (RuBi-glu, Tocris) (Fino *et al*., 2009) diluted into the bath solution for a final concentration of 800 µM as described (Tazerart *et al*., 2020). Following a 15-20 minutes resting period, cells were perfused with Rubi-glu. Monitoring changes in calcium at the spine head was done by imaging 2P calcium signals and detecting the fluorescence with two PMTs placed after wavelength filters (525/70 for green, 595/50 for red). The baseline spine head Ca^2+^ fluorescence signal was monitored for 200-400ms using low power 810 nm laser illumination on sample (∼8mW). Uncaging of RuBi-glu was performed at 810 nm (∼25–30 mW on sample) at a location adjacent to the selected spine head (≍0.2 µm away from the upper edge of the selected spine head, red dots in Fig.5), in the z-plane of maximal spine fluorescence intensity. Then, the line scan resumed for 800 ms at low imaging laser power (∼8mW on sample). This protocol was performed 10 times and the average ΔG/R was calculated from the corresponding signals as described (Tazerart *et al*., 2020) via a custom written Matlab script.

### Morphology

Individual spine morphology (estimated head volume, spine neck length) was assessed using a projection of a z-stack encompassing the spine of interest and the connecting segment of dendrite (*Fiji, NIH*). The spine neck length was measured from the proximal edge of the spine head to the edge of the dendrite or by computing the shortest orthogonal distance between the base of the spine head and the edge of the dendrite. This estimation was at times necessary because it was impossible to precisely ascertain the topology of the spine neck because of its small dimensions. For spines with no discernible necks, we chose a minimum value of 0.2 µm. Spine head diameter was measured by taking the average of the longest and corresponding orthogonal spine head dimensions. These were obtained by fitting a Gaussian curve to the profile of fluorescence across these axes, obtained from a z-stack encompassing the spine head (Δz = 0.4 µm). The corresponding spine head volume (V, in µm^3^) was estimated using the following formula:

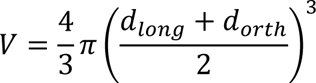

where dlong and dorth are the FWHM of the spine head along the longest and corresponding orthogonal dimension, respectively. The FWHM was calculated using the standard deviation (σ) of the Gaussian curve obtained from fitting the fluorescence intensity profile of the spine (FWHM = 2.335 σ).

### Statistics

Western blot densitometric quantification was analysed using a two-tailed paired t-test. Difference in dataset comprising two groups were measured using Mann-Witney test or Wilcoxon signed-rank test when appropriate. When more than two groups were measured, an ANOVA followed by a Tukey post hoc test was used. Statistical significance threshold was set at p<0.05*, p<0.01** and p<0.001***, p<0.0001****. Data are shown as mean ± standard error of the mean (S.E.M).

### Modelling

A morphologically realistic L5 PN NEURON model was used as a basis for this study (Nevian *et al*., 2007). Two 1 µm long spines were connected halfway along a 237 µm basal dendrite that was randomly selected from the basal arborization. Neuron simulations were run by sequentially varying spine head volume and BK conductance (gBK, see below). The neck width was estimated from the calculated head volume using published values (Arellano *et al*., 2007a). Neck and spine head axial resistances were set to 500 and 150 MΩ respectively (O’Donnell *et al*., 2011; Harnett *et al*., 2012). A high voltage-dependent Ca^2+^ channel (HVA, https://senselab.med.yale.edu/modeldb/ShowModel?model=135787&file=/ShuEtAl20062007/ca.mod#tabs-2) and Ca^2+^ buffering/removal mechanisms (resting concentration of 1nM (Destexhe *et al*., 1993)) were inserted in the spine head. Alpha subunits of BK channels (Jaffe *et al*., 2011)) were added in one spine (BK spine) while the other spine served as a control (gBK = 0 nS).

A physiologically realistic ionotropic glutamate receptor stimulation mechanism (Polsky *et al*., 2009) was added to each spine. The AMPA and NMDA conductance was calibrated to trigger physiologically relevant EPSPs at the soma (∼0.4−0.8 mV). The voltage-gated calcium channel conductance was then calibrated in order to reach physiologically relevant calcium changes in the spine head (sub 0.2 mM). During a simulation, the control spine was first activated. One second later, the spine with BK channels was activated. A stimulus run lasted 2 seconds. A series of 64 runs were obtained by iteratively varying the BK conductance (gBK from 0 to 30 nS in 2 nS increments) then head volume (0.065, 0.11, 0.18, 0.27 µm^3^). Runs were recorded in NEURON (Hines & Carnevale, 1997) as vectors and analyzed offline using Matlab.

## Results

### Expression of BK channel in visual cortex and L5 PN dendrites and spines

To detect the presence of the kcnma1 gene transcripts encoding the pore-forming α subunit (αBK) of BK channels in the mouse visual cortex, we analyzed the Allen mouse brain atlas *in-situ* hybridization data (Fig. 1A). The 522 riboprobe used for the detection of the kcnma1 gene has between 86-100 % homology with the 22 kcnma1 splice variants described in the mouse and thus is likely to recognize all kcnma1 splice variants. This data from the visual cortex indicates that kcnma1 is present in the soma of L2 to 6 neurons and that the majority of neurons, including the big somas of L5 PNs, are positive for the kcnma1 riboprobe (Fig. 1A).

**Figure 1.**
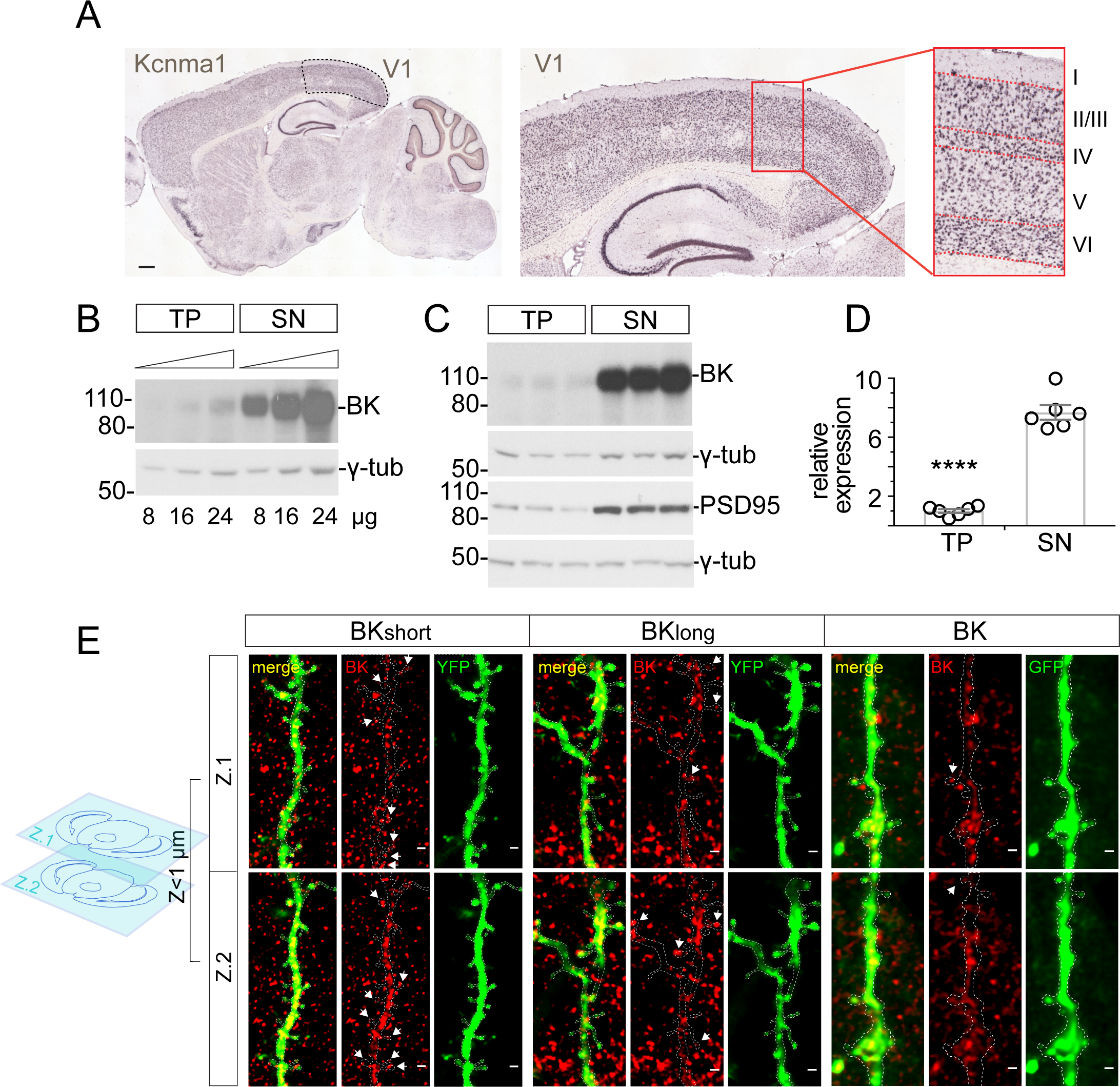
Expression of BK channel in the cortex and in the excitatory synapse: (A) Expression analysis of kcnma1 gene, encoding BK channel α subunit (αBK), in mouse cortex by in-situ hybridization shows expression in neuronal soma of cortical layers II/III to VI of the visual cortex area 1 (V1), where recordings in this study were made, Scale bar 500 μm. (B-D) Mouse visual cortex total protein (TP) and Synaptoneurosome (SN) fractions were analysed by Western Blot using a monoclonal αBK antibody and normalized by the expression of γ-tubulin. Synaptic enrichment in the SN fraction was confirmed by the expression of the post-synaptic density (PSD) marker PSD95. (B) A mix of the samples was loaded at increasing concentrations (8, 16, 24 ug/line) to confirm that the signal was not saturated and (C) loaded individually (16 ug/line) to access inter-individual variations. (D) Normalized BK expression in two independent experiments was plotted relative to the expression in the TP fraction (n = 6 p<0.0001). (E) Expression of BK channel in excitatory synapses was studied by immunofluorescence of αBK (red) in transgenic mice expressing YFP or GFP in L5 PN (green). Dendritic spine channel expression in each of two consecutive 0.7-0.9 μm confocal optical slices is indicated by arrows and seen in yellow (merge) when both signals are of similar intensity. Tissue expressing GFP (BK monoclonal antibody, right panels) was fixed in 4% PFA in PBS and tissue expressing YFP (BKshort and BKlong custom antibodies, right panels) corresponds to littermates of the mice used for electron microscopy, fixed at the same time in 4% PFA with 0.2% picric acid and 0.05% glutaraldehyde in 100 mM Sorensen’s phosphate buffer (pH 7.4) and processed using the same antibodies. Scale bar 1 μm.

Proteins are either translated from neuronal transcript in the soma and transported after synthesis to dendrites and axons (Hirokawa *et al*., 2010) or can be translated locally within axons (Jung *et al*., 2012) and dendrites (Bramham & Wells, 2007; Racca *et al*., 2010) themselves. We thus investigated the subcellular localization of BK channels in the cortex and found that this protein is highly enriched at excitatory synapses (p < 0.0001), as shown by Western blot analysis of synaptoneurosome (SN) fractions, obtained from a total protein (TP) homogenate from visual cortex (Fig. 1B-D). Care was taken to work in conditions where the signal was not saturated (Fig. 1B). Our results are in accordance with BK channel brain expression being predominant in axons and synaptic terminals (Trimmer, 2015). Since SN contain both pre- and post-synaptic membranes from excitatory synapses from a mixture of neurons, it is not possible to determine the precise cellular and subcellular localization of BK channels. To detect cell type-specific BK expression we used immunofluorescence on brain sections of transgenic mice expressing GFP or YFP in L5 PNs using two primary anti-BK channel antibodies detecting two splice variants (short and long, see *Material and Methods*) (Fig. 1E). We focused our attention on the basal dendrites of L5 PNs, where we performed the functional studies shown in Figures 4 and 5 and found that the BK channel is present in dendrites and dendritic spines with spine heads spanning different sizes (Fig. 1E). The expression of BK channels has been reported in dendrites throughout cortical layers of somatosensory cortex in rat brain (Benhassine & Berger, 2005). Therefore, the background fluorescence found in our analysis likely corresponds to expression of BK channels in other cortical neurons not expressing GFP or YFP. In addition, it was previously reported that BK channel expression in mouse hippocampal PNs is present in both pre and post-synaptic terminals (Sailer *et al*., 2006), although to our knowledge the subcellular profile of BK channels at the level of dendrites and spines in cortical pyramidal neurons or their presynaptic terminal has not been reported.

We then performed immunoelectron microscopy to detect the precise location of BK channels in dendritic spines located in the basal dendrites from L5 PNs with nanoscale resolution. To do so we used tissue from visual cortex expressing YFP in L5 PNs. We performed postembedding immunogold labeling for YFP and BK channels with gold particles of 12 and 6 nm, respectively. Our results showed that BK channels are located in spines. Interestingly, the gold particles labelling for BK channels were on the postsynaptic membrane both outside and inside of the post synaptic density (PSD), at the synaptic cleft and the spine neck (Fig. 2). Some gold particles were also observed in the cytoplasm (Fig. 2). In addition, we found examples of YFP-unlabeled PNs in L5 that also express BK channels in spines (Fig. 2E). In total we collected 35 labeled synaptic profiles, 15 corresponding to YFP-labeled L5 PNs, 11 from YFP-unlabeled PNs and 10 having positive signals for BK channels in the presynaptic sites. The PSD length measured in pooled 35 spines is 255.9 nm in average (rage from 121 to 415 nm), which is the typical synapse size in rodents (Kim & Sheng, 2009). The BK channel-expressing spine head volume was estimated as the fitted ellipsoid, and the head size ranged from 0.03 to 1.00 µm^3^. The signals for BK are observed in a wide variety of spine head sizes.

**Figure 2:**
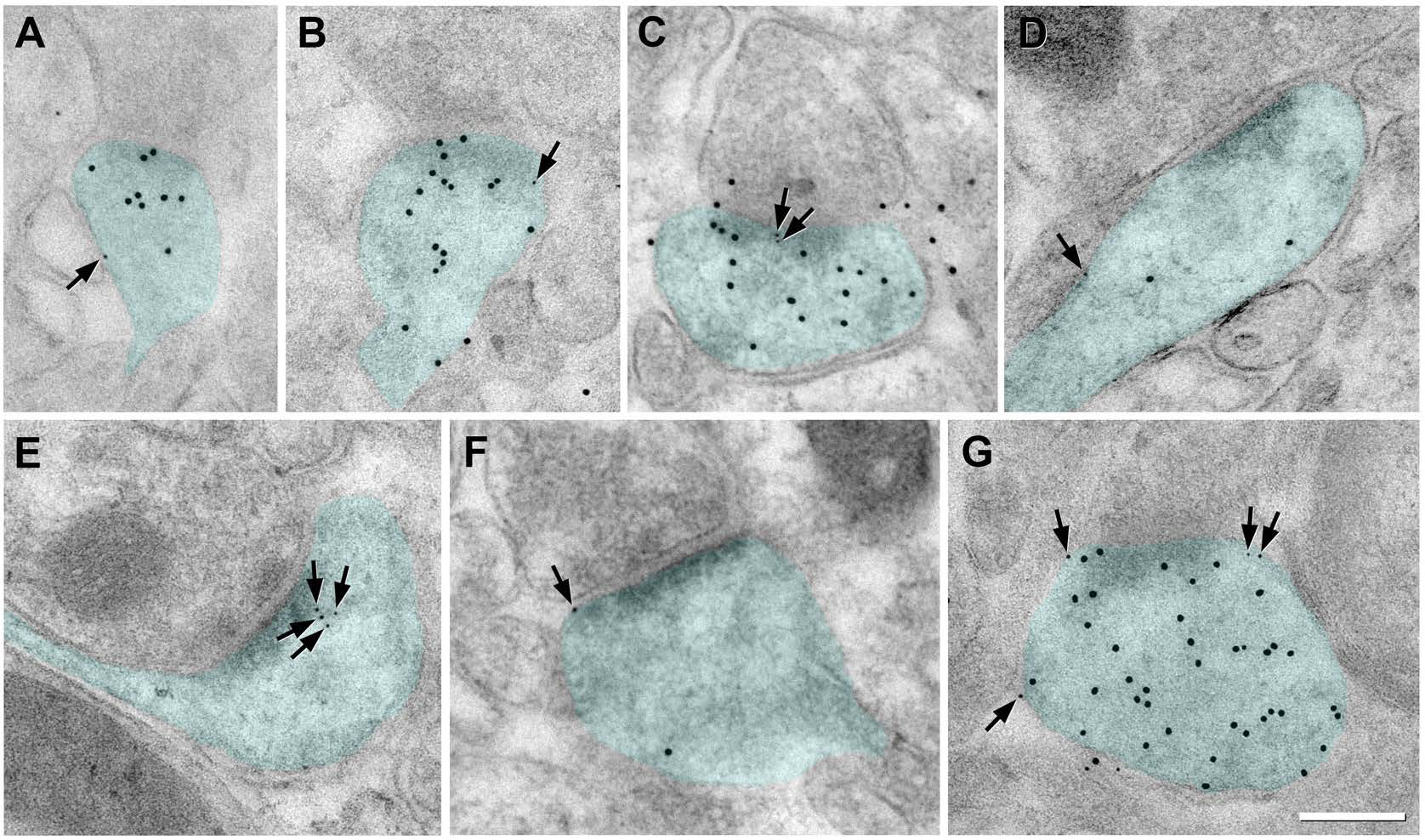
Localization of BK channels in spines visualized by immunoelectron microscopy. The signals for BK channels (6 nm immunogold particles, arrows) are located on membrane and in cytoplasm of spines of various sizes and shapes in both YFP expressing PNs (labeled with 12 nm gold particles, A-D, F, G) and unlabeled PN (E). Scale bar 200nm.

These results demonstrate with unprecedented resolution that BK channels are located in dendritic spines form basal dendrites of L5 PNs from visual cortex, and that this localization occurs in spines of different sizes and morphology (Fig. 2).

### Activation of BK channels in dendritic spines from basal dendrites of L5 PNs

Large conductance calcium-activated potassium (BK) channels are calcium- and voltage-activated, and dendritic spines can compartmentalize calcium (Yuste & Denk, 1995) and generate large voltage deflections at the spine head (Araya *et al*., 2007; Harnett *et al*., 2012; Araya, 2014). Hence, to test the hypothesis that functional BK channels located in dendritic spines modulate synaptic transmission we activated single spines using a method that mimics the synaptic release of glutamate in single dendritic spines via the 2P uncaging of caged glutamate (4-methoxy-7- nitroindolinyl glutamate (MNI)-glutamate, 2.5mM see *Material and Methods*) on dendritic spines located in the basal dendrites of L5 PNs from acute brain slices of mouse visual cortex as described (Araya *et al*., 2014; Mitchell *et al*., 2019; Tazerart *et al*., 2020) in the presence or absence of BK channel blockers or activators. Uncaging (u)-evoked excitatory postsynaptic potentials (uEPSP) were measured using a whole-cell patch clamp electrode in current-clamp configuration at the cell soma (Fig. 3A). This method allows for the selective activation of single dendritic spines with responses that are almost indistinguishable to the physiological activation of a single synapse ((Matsuzaki *et al*., 2001; Araya *et al*., 2006a; Araya *et al*., 2006b; Fino *et al*., 2009; Araya *et al*., 2013; Araya, 2014; Araya *et al*., 2014; Mitchell *et al*., 2019; Tazerart *et al*., 2019) and see *Material and Methods*). Briefly, we performed 2P uncaging of MNI-glutamate in spines under control conditions (10 uncaging repetitions at 0.5 Hz, in standard artificial cerebrospinal fluid (ACSF), see *Material and Methods*) and after bath application of Charybdotoxin (ChTx), a scorpion toxin that blocks BK channels with nM affinity (Miller *et al*., 1985a). Importantly, the repetitive 2P uncaging of MNI-glutamate to activate a single dendritic spine under control or after bath application of 100 nM of ChTx generates reliable uEPSP amplitudes without any trial to trial rundown (n = 10 uncaging events/activated spine) (-2.01 ± 1.22 % variation from initial amplitude in control, and -0.79 ± 1.09 % under ChTx, p = 0.39 Wilcoxon rank test, n = 27 spines, 16 neurons, 15 mice). Furthermore, bath application of 100 nM ChTx did not affect the passive membrane properties (*input resistance* in control: 131.7 ± 11.1 MΩ; or in ChTx: 123.2 ± 8.6 MΩ, n = 22 neurons, p = 0.17, Wilcoxon test, Fig. 3B left plot, the *leak current* in control: - 43.50 ± 8.7 pA; or in ChTx: -31.0 ± 6.7 pA, n = 30 neurons, p = 0.14, Wilcoxon test, Fig. 3B right plot). Bath application of ChTx induced a significant increase in AP width (control: 2.23 ± 0.15 ms; ChTx: 3.23 ± 0.32 ms, n = 12 neurons, p = 0.01, Mann-Whitney test Fig. 3C left traces and middle plot, n = 12 neurons) as previously shown in PNs from the amygdala (Faber & Sah, 2002), hippocampus (Lancaster & Nicoll, 1987) and in cortical L5 PNs (Benhassine & Berger, 2009; Bock & Stuart, 2016), and a significant increase of the AP decay time (control: 2.00 ± 0.17 ms; ChTx: 3.38 ± 0.43 ms, n = 12 neurons, p = 0.005, Mann-Whitney test, Fig. 3C right plot) without affecting the average inter-spike interval (ISI) or the initial ISI of the first two APs (p < 0.05, Mann-Whitney test, Fig. 3D and E).

**Figure 3.**
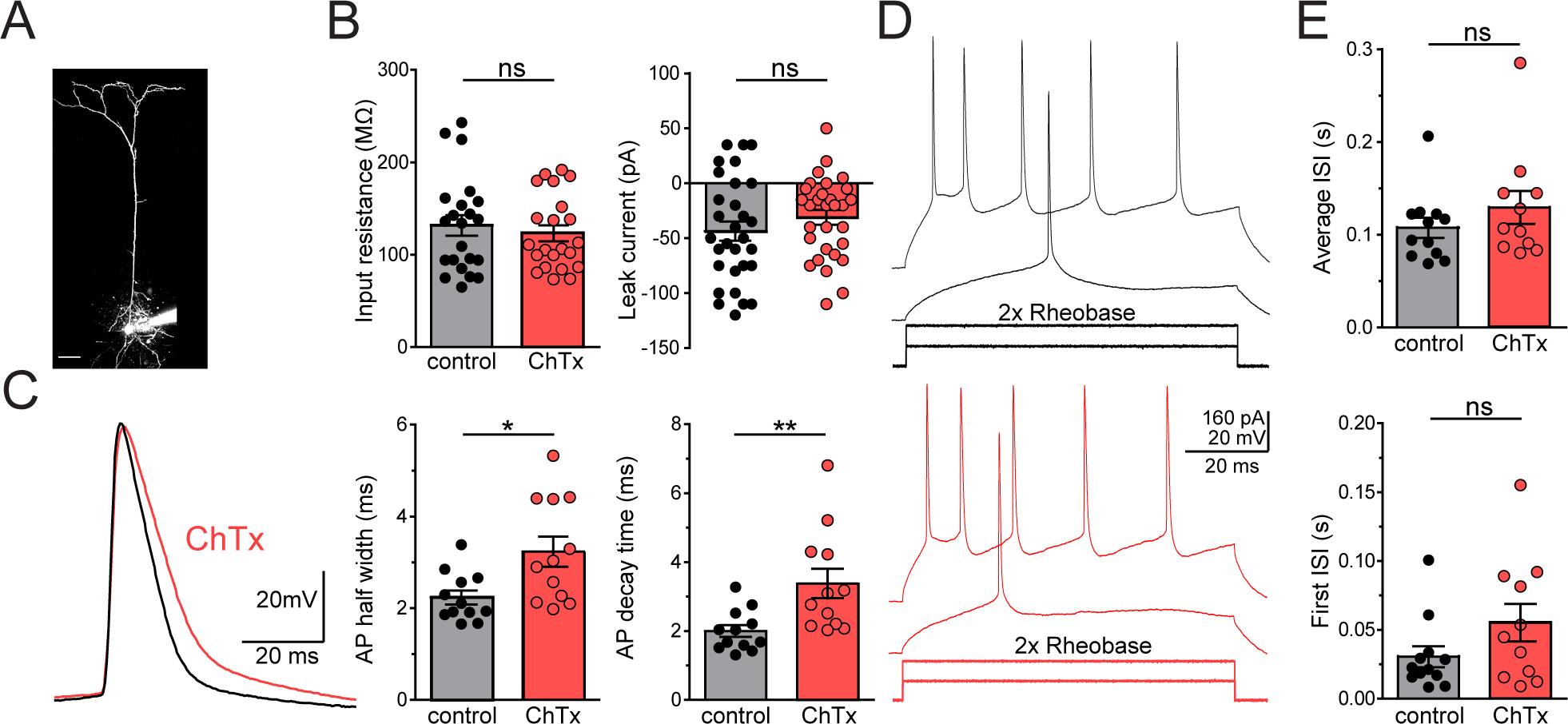
Effect of BK channels block on the electrophysiological properties of L5 PNs. (A) Two-photon (2P) image of a L5 PNs loaded through the patch electrode with Alexa 568 dye (120µM). (B) Input resistance (left) and leak current (right) of L5 PNs recorded in whole-cell patch-clamp configuration in control condition (black bars and dots) and in presence of the BK channels blocker (ChTx, 100nM, red bars and dots). (C) Left: Action potential (AP) examples recorded before (black trace) and after blocking BK channels (red trace) with ChTx. Average AP half-width (middle plot) and decay time (right plot) in control (black bars and dots) and in the presence of ChTx (red bars and dots). (D) Voltage responses during current injections at the soma (quadratic pulses) at once (middle trace) or twice the rheobase of the AP in control (black traces) and after blocking BK channels (red traces). (E) Average interspike interval (ISI, upper plot) and ISI of the first two AP (lower plot) measured at twice the rheobase in control (black bars and dots) and after blocking BK channels (red bars and dots). Scale bar on A panel = 50µm. * p<0.05, ** p<0.01, Mann- Witney test.

We then examined the uEPSP amplitude obtained in experiments after 2P activation of spines in control and in the same spines after bath application of 100 nM of ChTx (Fig. 4A, example of the 2P activation of a small-headed and a large-headed spine in the image above and the one below, respectively). Pooling all the data, we found that the average uEPSP amplitude was not significantly different between control and ChTx conditions (control: 0.59 ± 0.06 mV; ChTx: 0.6 ± 0.07 mV, n = 30 spines, 17 neurons, 16 mice, p = 0.9, Wilcoxon test, Fig. 4B). The morphology of the stimulated spines measured by 2P imaging was also unchanged in the presence of ChTx (neck length (NL) control: 1.21 ± 0.13 µm and 1.14 ± 0.13 µm under ChTx, p > 0.05, Wilcoxon rank test; head volume control: 0.14 ± 0.01 µm^3^ and 0.13 ± 0.01 µm^3^ under ChTx, p > 0.05, Wilcoxon rank test, n = 30 spines, 17 neurons, 16 mice). Since we found a significant variability in the effect of ChTx within the pool of activated spines, with some spines exhibiting a significant increase in the uEPSP amplitude after the addition of ChTx (Fig. 4B, left plot), we hypothesized that the spine head morphology ̶ which can affect the impedance and calcium accumulation in the spine head (Araya, 2014) ̶ is responsible for selectively activating spine BK channels. More specifically, we hypothesized that BK channels are predominantly activated in small-headed spines, which due to their higher impedance trigger large voltage deflections, and a large increase in glutamate-induced calcium responses at the spine head (Noguchi *et al*., 2005). Indeed, our data show that in spines with small head volumes there is an increase in the uEPSP amplitude following ChTx application (Fig. 4A). We then analyzed the ratio of uEPSP amplitude in the presence of ChTx to that of control conditions for all experiments and found that the ChTx-dependent increase in uEPSP amplitude decayed exponentially as a function of the spine head volume with a λ of 0.09 µm^3^ (Fig. 4B, right plot). Therefore, we used this value (λexp = 0.09 µm^3^) as the boundary to separate the 2P-activated spines into two groups: 1) small-headed spines (spine head volumes of ≤ 0.09 µm^3^) and 2) large-headed spines (> 0.09 µm^3^). By separating the data in this manner, we found that indeed the addition of ChTx induced a significant increase in the uEPSP amplitude of small-headed 2P-activated spines (≤ 0.09 µm^3^; 133.1 ± 11.86%, n = 10 spines, p = 0.021 different from control, Paired t-test, Fig. 4E), without significantly altering the uEPSP amplitude in large-headed spines with (> 0.09 µm^3^; 92.01 ± 5.22 %, n = 20, p = 0.14, Paired t-test, Fig. 4A, B and E). To further validate the functional contribution of BK channels in spines during synaptic transmission we performed 2P uncaging of glutamate in single spines in the presence or absence of the specific synthetic BK activator NS1619 (1-(2’-hydoxy-5’-trifluoromethylphenyl)-5- trifluoromethyl-2(3H)benzimidazolone). This synthetic benzimidazolone derivative reversibly triggers a dose-dependent shift in the voltage-activation curve of the channel to more negative potentials, hence, increasing the open probability of BK channels without affecting single channel conductance properties (Olesen *et al*., 1994).

**Figure 4.**
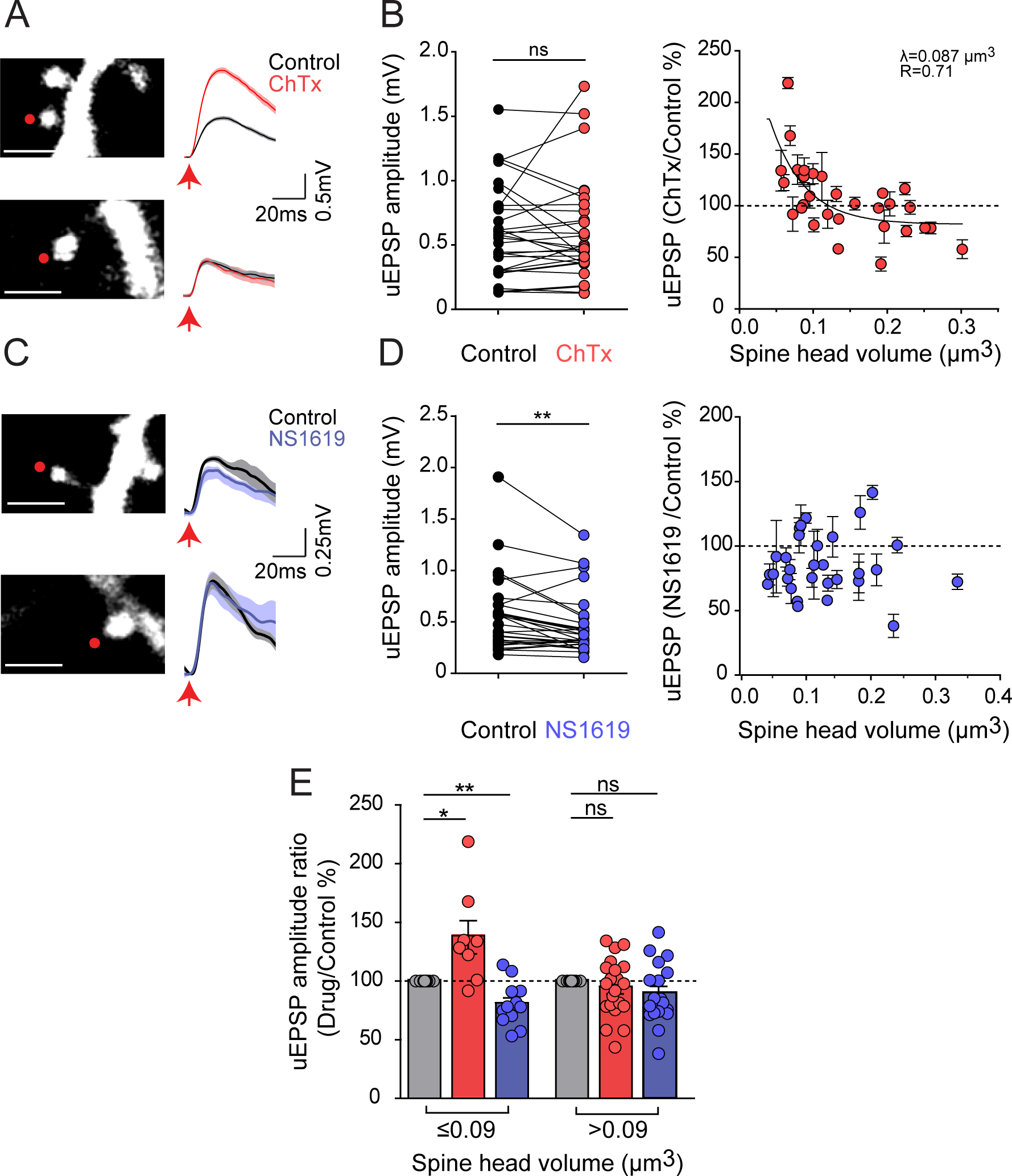
Selective activation of BK channels in small-headed dendritic spines suppress excitatory postsynaptic potentials. (A) Representative experiments where a spine was selected for 2P glutamate uncaging stimulation in the absence and after the addition of the BK channel blocker ChTx. Traces correspond to average of ten uEPSPs recorded in the soma and generated by 2P uncaging before (control, black trace) and after adding ChTx (100nM, red trace) over the indicated spine (red dot). (B) Left, uEPSP amplitude before (black dots) and after adding ChTx 100nM (red dots). Right: plot of the ratio of uEPSP amplitude in the presence of ChTx to that in control conditions as a function of spine head volume of the activated spines. Data was fitted with an exponential (see *Material and Methods* section) and λ calculated. (C) Representative experiments where a spine was selected for 2P glutamate uncaging stimulation in the absence and after the addition of the BK channel activator NS1619. Traces correspond to average of ten uEPSPs recorded in the soma and generated by 2P uncaging before (control, black trace) and after adding NS1619 (100nM, blue traces). (D) Left, uEPSP amplitude before (black dots) and after adding NS1619 (100nM, blue dots). Right: plot of the ratio of uEPSP amplitude in the presence of NS1619 to that in control conditions as a function of spine head volume of the activated spines. (E) Changes in the uEPSP amplitude ratio (drug/control%) for the BK blocker ChTx (red dots and bars) and activator NS1619 (blue dots and bars) in small-headed (volume ≤0.09µm^3^) and large-headed spines (> 0.09 µm^3^). Scale bars on A and C = 2µm. * p<0.05, **, p<0.01, Wilcoxon test in B and D and paired t-test in E.

Our results showed that bath application of 100 nM NS1619 does not affect the input resistance of neurons (control: 163.2 ± 12.02 MΩ,; NS1619: 160.2 ± 13.77 MΩ, n = 13 neurons, p = 0.73, Wilcoxon test), the leak current (control: -45.5 ± 9.98 pA; NS1619: -47.07 ± 13.37 pA, n = 30 neurons, p = 0.62) the AP half width (FWHM of AP in control: 2.42 ±0.13 ms; NS1619: 2.38 ± 0.11, p = 0.62, Wilcoxon ranked test, n = 16 neurons) and the AP decay time (control: 2.24 ± 0.11 ms; NS1619: 2.24 ± 0.11 ms, p = 0.76, Wilcoxon ranked test, n = 15 neurons).

We then analyzed the effect of bath application of NS1619 on the uEPSP of the 2P-activated spines. Pooled data showed that bath application of NS1619 resulted in a significant reduction of the uEPSP amplitude (control: 0.56 ± 0.07 mV; NS1619: 0.45 ± 0.05 mV, n = 30 spines, p = 0.0018, Wilcoxon test, Fig. 4D), as expected by the increased BK channel open probability actions of NS1619 (Olesen *et al*., 1994). However, we found significant variability in the effect of NS1619 within the pool of activated spines, with some spines exhibiting a significant decrease in the uEPSP amplitude after the addition of NS1619, while no effect in the uEPSP amplitude was evident in others (Fig. 4C and D). Since the spine head volume is an important factor in determining the activation of BK channels (Fig. 4A and B) we hypothesized that the BK channel enhancement actions of NS1619 is preferentially observed in small-head spines. Indeed, addition of NS1619 significantly reduces the uEPSP amplitude in small-headed spines (≤ 0.09 µm^3^ head volume, n = 12, p = 0.004, paired t-test Fig. 4C upper image and traces and Fig. 4E), but not in large-headed spines – although there is a non-significant tendency for a reduction in uEPSP amplitude (> 0.09 µm^3^, n = 18 spines, p = 0.1, Fig. 4C lower image and traces and Fig. 4E). Taken together these results suggest that although BK channels are expressed in spines of different morphologies (Figs. 1-2), they can only be activated in small-headed spines during synaptic transmission in L5 PNs, reducing synaptic weight of those synapses at the soma (Fig. 4).

Since BK channels are voltage- and calcium-activated potassium channels and given the observed spine head volume dependence with BK channel activation, we next performed experiments to directly measure calcium signals at the spine head of activated spines. To do this, we performed nearly simultaneous 2P uncaging of caged glutamate in single spines and 2P imaging of calcium in the activated spine head with high spatiotemporal precision, using the linescan imaging mode from a line crossing the activated spine head (Fig. 5A) as described (Mitchell *et al*., 2019; Tazerart *et al*., 2020), while recording the uEPSPs at the soma (Fig. 5A and B). Changes in spine head calcium concentration were measured using the low-affinity calcium sensitive dye Fluo-4 (He *et al*., 2014) and Alexa Fluor 594 in L5 PNs held at -65 mV using a somatic current clamp electrode. Fluorescence was computed as the relative change in calcium concentration ΔG/R as described (Tazerart *et al*., 2020). Pooled data show that the average change in fluorescence of the calcium indicator from baseline following 10 uncaging events at each activated spine varies among spines with different morphologies (peak calcium amplitude of 108.4% ± 0.52% ΔG/R, ranged between 102.7% and 119% ΔG/R, n = 48 spines). These spine head calcium signal amplitudes are consistent with previous results using similar methods (Sabatini *et al*., 2002; Bloodgood & Sabatini, 2007). We next investigated whether there was a correlation between the head volume of the 2P activated spines and the 2P calcium responses at the spine head. We found a significant negative correlation (p = 0.02), whereby calcium signals in small headed spines were larger than those observed in big headed spines (Fig. 5B). Then, we binned our data based on the λexp obtained from the exponential fit from the ChTx-dependent increase in uEPSP amplitude as a function of the spine head volume (Fig. 4B) to separate the data into small-headed (≤ 0.09 µm^3^) and large-headed dendritic spines (> 0.09µm^3^) groups. We found that small-headed spines had significantly higher calcium responses in the spine head that those of large-headed spines (Fig. 5B, right plot; Spine head volume ≤ 0.09 µm^3^: 110.5 ± 0.97 %ΔG/R, head volume > 0.09 µm^3^: 107.2 ± 0.51% ΔG/R, p = 0.002, Mann-Whitney test). Furthermore, we found a significant correlation (p = 0.02) between the spine head volume and the uEPSP recored at the soma (Fig. 5B inset).

**Figure 5:**
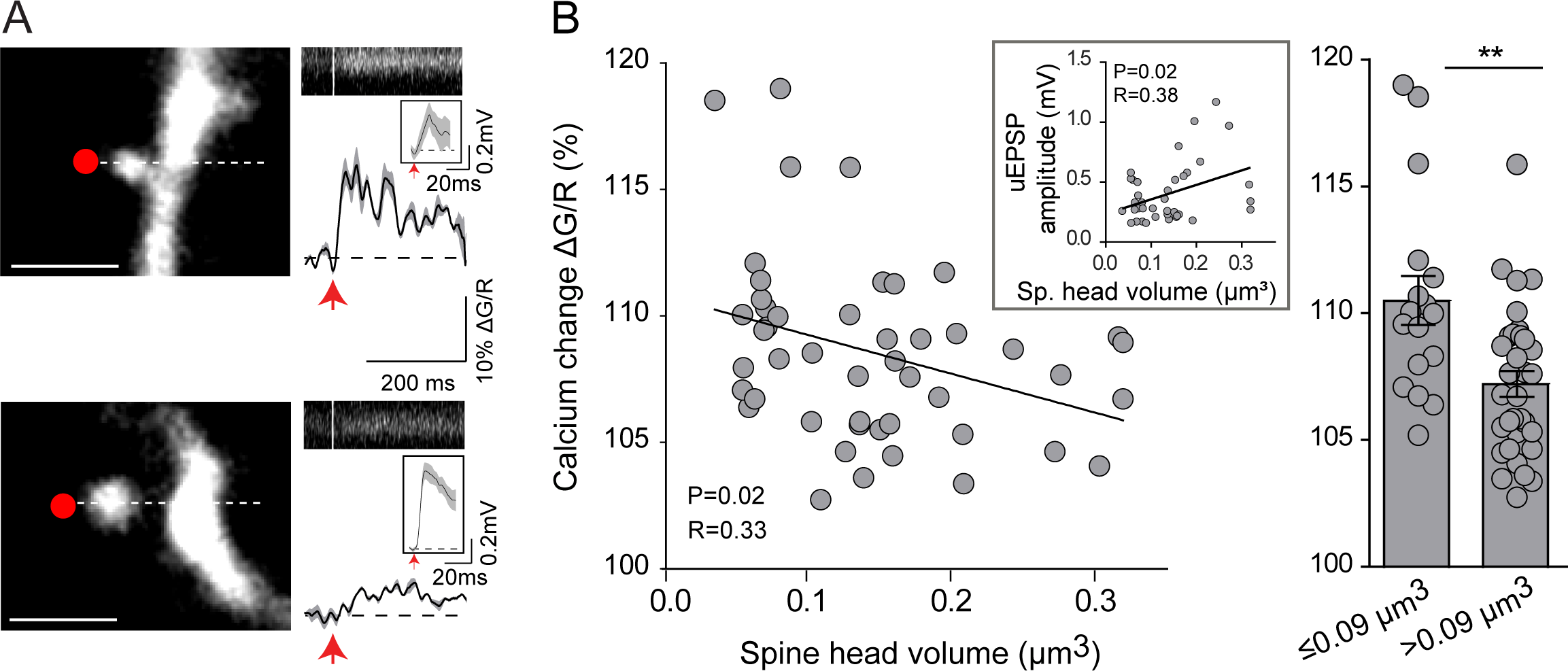
Inverse correlation between spine head calcium responses and spine head volume. (A) Left images, representative experiments of 2P calcium imaging during 2P uncaging of spines with different spine head volumes. L5 PNs were loaded with Alexa 594 (100 µM) and a calcium- sensitive dye (Fluo-4, 300 µM). White dotted lines depict the position of the 2P imaging line-scan (see *Material and Methods* section) while the red dots indicate the location of the laser beam during the uncaging pulse (see *Material and Methods* section). Right, traces correspond to the average of ten calcium responses (Δ G/R, black traces) at the spine head of the 2P uncaging-activated spines, and insets the average uEPSP of the 2P activated spine. The image above the calcium and uEPSP traces is the 2P-calcium fluorescence over time from the region intersected by the dotted line in left images. (B) Left, the inverse correlation between 2P-calcium signals at the spine head and spine head volume of the 2P uncaging-activated spines. Inset, the correlation between the uEPSP amplitude recorded at the soma and the spine head volume of the activated spines. Right, a plot showing calcium changes at the spine head for small-headed spines (≤ 0.09µm^3^) and large-headed spines (> 0.09 µm^3^). The shaded gray area in the calcium traces from A shows the SEM. ** p<0.01, Mann-Whitney test. Scale bars in A = 2µm.

Hence, our results indicate that small-headed dendritic spines (≤ 0.09 µm^3^) generate larger 2P- induced calcium responses than those observed in spines with bigger head volumes during synaptic transmission and selectively activate BK channels to suppress uEPSPs.

### Biophysical simulations of BK channel activation in dendritic spines

We then explored the selective activation of BK channels in small-headed spines using multicompartmental simulations in the NEURON environment (Hines & Carnevale, 1997). We simulated our experiments by building a morphologically realistic L5 PN model (adapted from (Nevian *et al*., 2007)) to assess the putative role of spine BK channels in synaptic transmission (Fig. 6). Briefly, we built a spine with a 1 µm long neck and a head volume that ranged from 0.065 - 0.3 µm^3^ to cover the range of spine head volumes probed in our 2P experiments. The dendritic spine was connected to a basal dendrite of the modeled L5 PN (140 µm away from soma) and AMPA-receptors(R), NMDA-R, voltage-gated calcium channels (VGCC) and BK channels were placed in the plasma membrane of the spine head compartment (see *Material and Methods*) (Fig. 6A1 and B1). We simulated synaptic responses and EPSPs with amplitudes similar to those recorded in L5 PNs under physiological conditions or after the 2P uncaging of glutamate in single spines (Araya *et al*., 2014). EPSP amplitudes recorded at the soma showed a positive correlation with spine head size similar to what we observed in our experiments, with EPSPs increasing from 0.45 to 0.8 mV as spine head volume increased from 0.065 to 0.27µm^3^ (Fig. 6A2 and B2, and C inset). We then explored the contribution of BK channels to EPSP amplitude by adding BK channels to spines (ranging from 0-30 nS, or ∼ 0 to 120 channels assuming unitary BK channel conductances of ∼ 250 pS), and found that the activation of spine BK channels reduced the EPSP amplitude recorded at the soma in a manner that strongly depended on the head volume of the activated spine (Fig. 6A2, B2 and C1 inset) similarly to what we observed in our experiments (compare Fig. 4 and 6). More specifically, the effect of blocking BK channels on somatic EPSP amplitude (calculated as the somatic EPSP amplitude in absence of BK channels (gBK = 0nS) divided by the somatic EPSP amplitude in the presence of different BK conductances) was modulated in small-headed spines without altering large-headed spines (Fig. 6C1 and C2). These data fit with an exponential decay resulted in a modeling λ (λmod) ranging from 0.057 µm^3^ when gBK is set at 2nS to 0.099 µm^3^ when gBK is at 30 nS (Fig. 6C1). For the range of gBK values modeled, the calculated λmod is similar to our λexp, but is nearly identical when gBK is set at 14nS, representing ∼56 BK channels per spine.

**Figure 6:**
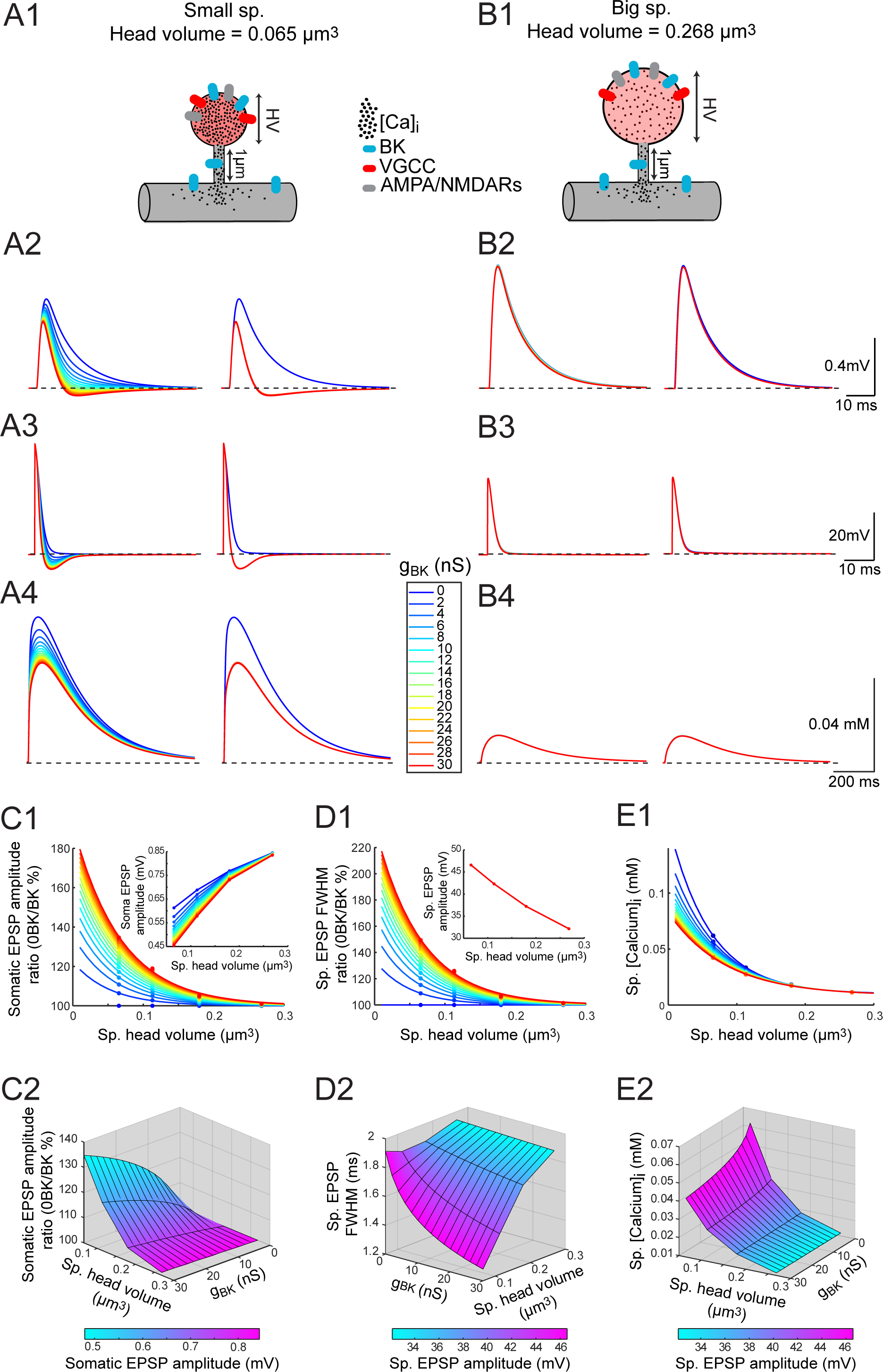
Biophysical modeling of the activation of BK channels in dendritic spines of varying head volumes and their role in synaptic transmission. (A1 and B1) Schematic of the modeling approach in small-headed (A1) and large-headed spines (B1) expressing BK channels, AMPA receptors (AMPAR), NMDAR, and voltage-gated calcium channels (VGCC). (A2 and B2) Measurements of EPSPs at the soma in the presence of varying spine BK channel conductances (0-30 nS). Note how in small-headed spines (A2), but not in large-headed spines (B2), there is significant activation of BK channels that reduces somatic EPSP amplitude. (A3 and B3) Measurements of EPSPs at the spine head in the presence of varying spine BK channel conductances (0-30 nS). Note how in small-headed spines (A3), but not in large-headed spines (B3), there is a significant activation of BK channels that reduces the duration of spine EPSP. (A4 and B4) Measurements of calcium concentrations at the spine head in the presence of varying spine BK channel conductances (0-30 nS). Note how in small-headed spines (A4) there are larger calcium responses triggered than in large-headed spines (B4), and that the activation of BK channels reduces the amplitude of the spine calcium response only in small-headed spines (A4). A1-A4 shows data for a spine with a head volume = 0.065 µm^3^. B1-B4 is the same as in A1-A4 but for a spine with head volume of = 0.27 µm^3^. (C1) Inverse correlation between the effect of blocking BK channels on EPSP amplitude (*somatic EPSP amplitude ratio*; calculated as the somatic EPSP amplitude in absence of BK channels (gBK = 0nS) divided by the somatic EPSP amplitude in the presence of different BK conductances) and spine head volume modeled with different BK conductances. Note that the BK-dependent effects on somatic EPSP amplitude, resulting in a decrease in somatic EPSP amplitude (inset), only occurs in small-headed spines. (C2) Relationship between somatic EPSP amplitude ratio, spine head volume, BK conductance and somatic EPSP amplitude. (D1) Plot illustrating the effect of blocking BK channels on spine EPSP FWHM (*spine EPSP FWHM ratio*; calculated as the spine EPSP FWHM of the activated spines with no BK channels (gBK = 0nS) divided by the spine FWHM duration in the presence of different BK conductances) and spine head volume modeled with different spine BK conductances. Inset shows the relationship between the spine head EPSP amplitude and the spine head volume. Note how the spine EPSP amplitude is not affected in small-headed spines where BK channels are activated. (D2) Relationship between spine EPSP FWHM ratio, spine head volume, BK conductance and spine head EPSP amplitude. (E1) Inverse correlation between the spine head calcium concentration and the spine head volume modeled with different spine BK channel conductances. (E2) Relationship between spine head calcium concentration, spine head volume, BK conductance and spine EPSP amplitude. Sp = Spine; gBK = conductance of BK channels.

We then explored EPSP amplitudes recorded at the spine head and found that there is a negative correlation with the spine head size, with fast spine head EPSPs of less than 10ms duration, decreasing from 47 to 32 mV as spine head volume increased from 0.065 to 0.27µm^3^ (Fig. 6A3 and B3, and D1 inset). We tested how BK channels shape EPSPs amplitude and duration directly at the spine head and found that the activation of spine BK occurs only in small-headed spines, reducing the duration of EPSPs without affecting the spine EPSP amplitude (Fig. 6A3, B3, D1 and D2). The BK-dependent reduction in spine head EPSP duration strongly depends on the head volume of the activated spine (Fig. 6A3, B3, D1 and D2). More specifically, the effect of blocking BK channels on the spine head EPSP duration, calculated as the spine EPSP full width at half maximum (FWHM) of the activated spines with no BK channels (gBK = 0nS) divided by the spine FWHM duration in the presence of different BK conductances, was increased in small-headed spines but not large-headed spines (Fig. 6D1 and D2). We then explored the corresponding head calcium concentration of the activated spines (Fig. 6A4, B4, E1 and E2) and found a spine head *volumetric* dependency on calcium concentration at the spine head. This dependency fit an exponential decay, where calcium concentrations decay exponentially as a function of spine head volume (Fig. 6E1). These modeling results closely matched our experimental results (Fig. 5B) and together indicate that BK channels when present in spines of different shapes can only be activated in small-headed spines, where large voltage and calcium concentrations are reached.

## Discussion

In this study we show that large conductance calcium-activated potassium (BK) channels are present in dendritic spines of all shapes in basal dendrites from L5 PNs, but they are selectively activated in small-headed spines (≤ 0.09 µm^3^, λexp = 0.09) and suppress uEPSPs. Using 2P uncaging of glutamate, we demonstrated that (1) bath application of Charbydotoxin (ChTx), a potent BK channel inhibitor (Miller *et al*., 1985a), does not affect uEPSP amplitude when considering all activated spines, but significantly increased the amplitude of uEPSPs only in small-headed spines (≤ 0.09 µm^3^); (2) bath application of the BK channel activator NS1619, a synthetic benzimidazolone derivative, significantly reduced the amplitude of uEPSP only in small-headed spines (≤ 0.09 um^3^); (3) intracellular calcium responses in the head of 2P-activated spines are significantly larger in small-headed spines (≤ 0.09 µm^3^) than those observed in spines with larger head volume; and finally, (4) using multicompartmental modeling of spines of varying head volumes (spanning the values obtained in our experimental dataset) we showed that during synaptic transmission BK channels are only activated in small-headed spines, where larger intracellular calcium concentrations and EPSPs amplitudes are generated in the spine head in contrast to spines with bigger head volumes.

Our results and previous published observations have shown that blocking somatic BK channels in L5 PNs significantly increased action potential (AP) width, without affecting firing frequency or the passive membrane properties (Benhassine & Berger, 2009; Bock & Stuart, 2016). Furthermore, local blockade of BK channel in the apical dendrites of L5 PNs has been shown to increase the duration of dendritic calcium spikes without affecting backpropagating (b)APs (Bock & Stuart, 2016) (although see (Benhassine & Berger, 2009)). Interestingly, these published observations showed that the amplitudes of bAPs, recorded 600 to 800 um away from the soma, triggered depolarizations of ∼35 mV to 15mV, respectively, that were insufficient to activate BK channels (Bock & Stuart, 2016) a result that it is consistent with other observations in hippocampal CA1 PNs (Poolos & Johnston, 1999). These results suggest that BK channels are not recruited at sub-threshold membrane potentials in cellular compartments, where insufficient voltage and calcium signals are generated. Based on these observations we wondered if dendritic spines, with their small size (< 1 fL volume) and high impedance, could generate local depolarizations and calcium signals at the spine head that would be sufficient to activate BK channels during synaptic transmission? Theoretical and experimental studies have suggested that the electrical function of spines during synaptic transmission is to generate a large EPSP at the spine head (Jack *et al*., 1975; Segev & Rall, 1988; Harnett *et al*., 2012; Araya *et al*., 2014) capable of activating spine voltage-gated ion channels at the spine but not in the parent dendrite. Activation of these channels has been shown to be an effective mechanisms for shaping synaptic weight (Miller *et al*., 1985b; Perkel & Perkel, 1985; Shepherd *et al*., 1985; Segev & Rall, 1988; Araya *et al*., 2007; Bloodgood & Sabatini, 2007). Experimental observations aimed at estimating the electrical properties of spines suggest neck resistances of ∼ 500 MΩ (Harnett *et al*., 2012; Araya *et al*., 2014; Jayant *et al*., 2017) (although see (Popovic *et al*., 2015)), which are sufficient to amplify spine potentials at the spine head to >20 mV (Harnett *et al*., 2012). In fact, direct measurements of EPSPs at the spine head showed amplitudes that fluctuate from ∼10 to more than 50 mV at the spine head (mean 26 mV) using quantum-dot-coated nanopipettes (Jayant *et al*., 2017), similar to the range of voltages observed in our spine modeling results (Fig. 6D inset).

Spines can also act as biochemical compartments (Yuste & Denk, 1995), and theoretical and experimental data has indicated that spine neck and head morphology are important determinants on the decay time and amplitude of intracellular calcium signals in the spine head (for review see (Araya, 2014)). In fact, in hippocampal CA1 PNs it has been suggested that the spine head volume is negatively correlated with the amplitude of 2P-glutamate uncaging-generated spine calcium signal, but positively correlated with the intracellular calcium signals in the adjacent dendritic shaft (Noguchi *et al*., 2005). Similarly, our data shows experimentally that in L5 PNs small-headed spines (≤ 0.09 µm^3^) have larger intracellular calcium concentration than those observed in spines with bigger head volumes (> 0.09 µm^3^), following a negative correlation between the spine head volume and the 2P glutamate uncaging-generated spine intracellular calcium (Fig. 5). Importantly, our modeling results corroborate these findings, by clearly showing an inverse correlation between the spine head volume and the 1) intracellular spine head calcium concentration and 2) EPSP amplitude at the spine head, leading to increases in the open probability of BK channels (via its modular voltage sensors and calcium binding sites) only in small-headed spines (Latorre *et al*., 1989; Horrigan & Aldrich, 2002; Latorre *et al*., 2017). Hence, small-headed spines are perfectly poised to activate BK channels and to locally repolarize EPSPs.

What are the actual voltages and intracellular calcium concentration generated in small-headed spines during synaptic transmission? Our modeling results showed that the amplitude of the spine head voltages generated in small-headed spines capable to activate BK channels are > 40 mV, with intracellular spine head calcium concentration of > 27 µM, which are in agreement with previous estimations of the range of spine head intracellular calcium concentrations (Higley & Sabatini, 2012) and EPSP amplitudes (Harnett *et al*., 2012; Araya *et al*., 2014; Jayant *et al*., 2017), as well as with observations showing that BK channels are not activated by bAP in distal dendrites with recorded amplitudes of ∼ 35 mV (Bock & Stuart, 2016). Taken into account these published observations and our results, we can conclude that the activation of BK channels in L5 PNs occurs 1) during synaptic transmission only in small-headed spines to repolarize EPSPs; 2) during the induction of dendritic calcium spike in distal dendrites to control its duration (Bock & Stuart, 2016); and 3) during AP generation at the axon initial segment and soma widening AP amplitude (Benhassine & Berger, 2009; Bock & Stuart, 2016).

Dendritic spines are the minimal functional units of plasticity (Matsuzaki *et al*., 2001; Matsuzaki *et al*., 2004; Harvey *et al*., 2008; Nishiyama & Yasuda, 2015; Tazerart *et al*., 2020), and the induction of LTP at spines is thought to be the underlying mechanisms for learning and memory in the brain (Hayashi-Takagi *et al*., 2015). It has been reported that the spine head volume is an important factor in determining the induction of long-term potentiation (LTP) (Matsuzaki *et al*., 2004; Tazerart *et al*., 2020). More specifically, spines with head volumes of less than < 0.1 µm^3^ , which represent the group of spines capable of generating large voltage and intracellular calcium concentrations required to activate BK channels in mouse L5 PNs, are the preferential sites for the induction of LTP (Matsuzaki *et al*., 2004) and timing-dependent LTP (t-LTP) (Tazerart *et al*., 2020). Interestingly, the induction of LTP in spines triggers structural remodeling (neck shrinkage for early-LTP and spine volume enlargements for late-LTP) that is tightly coupled with synaptic function (Matsuzaki *et al*., 2004; Harvey *et al*., 2008; Araya, 2014; Araya *et al*., 2014; Tazerart *et al*., 2020). Furthermore, it has recently been suggested that BK channels affect the threshold for the induction of LTP in L5 PNs (Gomez *et al*., 2021). Hence, we hypothesize that one important *raison d’être* for the presence of BK channels in spines is to protect small-headed spines from promiscuously triggering LTP, keeping the reservoir of plasticity-prone spines from storing meaningless information but available for storing strong and persistent meaningful synaptic inputs. This BK channel-dependent control of plasticity could also apply to spike-timing-dependent plasticity (STDP), where we have recently shown that the single spine STDP rule has a narrow t-LTD and t-LTP induction time window (Tazerart *et al*., 2020). The narrow pre-post (for the induction of t-LTP) and post-pre (for the induction of LTD) timing for the single spine STDP rule could likely be finely-tuned by the BK channel-dependent shaping of EPSPs, generating a more precise and stringent gate for the induction of t-LTP and or t-LTD.

Interestingly, humanization of mouse cortical PNs in vivo via the expression of human-specific paralogs of SRGAP2A, SRGAP2C, induces neoteny during spine maturation (Charrier *et al*., 2012), increased cortico-cortical feedback and feedforward connectivity, reliability of sensory evoked responses, and improved associative learning tasks (Schmidt *et al*., 2020). Importantly, in juvenile mice humanization by SRGAP2C-expression in cortical PNs leads to the majority of spines having longer necks and smaller heads than their control counterparts (Charrier *et al*., 2012). These humanization-dependent spine structural changes are likely to increase spine impedance and with that the amplitude of EPSPs at the spine head, the intracellular calcium concentrations and ultimately the activation of spine BK channels in the majority of spines. Hence, we hypothesize that the repolarization of spine EPSPs by BK channels in humanized PNs occurs in the majority of spines and would lead to a more stringent and precise representation, storage, and associations of sensory and predictive synaptic inputs in PNs. These synaptic transmission modifications could be at the core of the improvements in associative learning tasks observed in these humanized PNs (Schmidt *et al*., 2020).

The degree of depolarization and intracellular calcium in spines has been shown to be influenced by neuromodulation (Giessel & Sabatini, 2010; Araya *et al*., 2013; He *et al*., 2014). In hippocampal CA1 PNs, M1 muscarinic receptor activation leads to higher synaptic potentials and calcium influx in spines via the inhibition of spine SK channels (Giessel & Sabatini, 2010). In addition, in cartwheel cells of the dorsal cochlear nucleus it has been shown that cholinergic signaling boost spine calcium responses and somatic EPSPs via L-type calcium channel-dependent blockade of BK channels (He *et al*., 2014). Hence, it is possible that the activation of BK channels in small-headed spines of L5 PNs could be modulated (i.e. blocked) by cholinergic signaling, leading to changes in synaptic weight. Future experiments are required to test this possibility and how it might affect synaptic transmission and synaptic plasticity.

Dopamine, another neuromodulator implicated in reinforcement effects on neuronal circuits in prefrontal cortex, has been shown to trigger calcium signals in the activated spine heads (Araya *et al*., 2013), and contribute to structural spine plasticity (Yagishita *et al*., 2014; Urakubo *et al*., 2020). Hence, one possibility is that dopaminergic signaling in spines from L5 PNs might influence glutamatergic transmission and the activation of BK channels in spines that otherwise would not be capable of activating these channels.

Novel experimental workflows are starting to allow us to reliably characterize the ultrastructure in functionally imaged neurons, and in particular to obtain a large number of ultra-structurally reconstructed spines and correlate them back to its initial 2P *in vivo* calcium imaging (Scholl *et al*., 2021; Thomas *et al*., 2021). Hence, the combination of *in vitro* or *in vivo* light-microscopy functional imaging and activation of spines with nanometer resolution features obtained with serial-block scanning electron microscopy (SBF-SEM) (Scholl *et al*., 2021; Thomas *et al*., 2021) are starting to pave the way for understanding the structural-functional passive and active spines properties. These approaches combined with 2P photoactivation of single synapses are shedding light into spine function during synaptic transmission, plasticity and integration of synaptic inputs. These technical breakthroughs reducing the timeframe and scope of the functional and structural probing of spines will likely help us in better understanding PNs and cortical function.

## Acknowledgments

We thank Alvaro Barrios for technical assistance and are grateful to all other members of Roberto Araya’s laboratory for kind support. We thank Debbie Guerrero-Given for technical support with EM sample preparation. We thank P. Drapeau for critical discussion and reading of the manuscript. Confocal microscopy experiments carried out at the Platform of Imaging by Microscopy (PIM) of the CHU Sainte-Justine Research Center (CHUSJRC) where supported by the expertise of E. Küster-Schöck. This work was funded by the Canadian Institutes of Health Research (CIHR) grant MOP-133711 to R.A., a Canada Foundation for Innovation (CFI) equipment grant Fonds des leaders 29970 to R.A., and a Natural Sciences and Engineering Research Council of Canada (NSERC Discovery Grant) grant application no. 418113-2012 (NSERC PIN 392027) to R.A. S.T. was supported in part by a salary support from the GRSNC at Université of Montréal. M.B. was supported by a Herber Jesper postdoctoral fellowship at Université of Montréal. D.E.M. was supported in part by a postdoctoral fellowship from the Fonds de recherche du Québec Santé (FRQS).

## Contributions

R.A. conceived the project. S.M.-R., performed the experiments and data analyses presented in Figure 1. C.I.T., S.M.-R., and N.K. performed the experiments and data analyses presented in Figure 2. S.T., and M.G.B., performed the experiments and data analyses presented in figures 3 and 4. S.T., performed the experiments and data analyses presented in Figure 5. D.E.M., M.G.B., and B.N-P performed the experiments and data analyses presented in Figure 6. R.A., M.G.B., S.T., C.I.T., N.K. and S.M.-R. designed experiments. R.A., S.T., M.G.B., S.M.-R., and D.E.M wrote the manuscript. R.A. supervised the project. All authors read and approved the contents of the manuscript.

